# Stochastic models of the growth dynamics of some dendritic cells

**DOI:** 10.1101/454264

**Authors:** Khairia El-Said El-Nadi Khairia

## Abstract

Different models of tumor growth are considered. Some mathematical methods are developed to analyze the dynamics of mutations enabling cells in cancer patients to metas-tize. The mathematical models consist of some stochastic dynamical systems describing tumor cells and immune effectors. It is also considered a method to find the ideal outcome of some treatments. Some different types of dendritic cells are considered. The obtained results will help to find some suitable treatments,which can be successful in returning an aggressive tumor to its passive,non-immune evading state. The principle goal of this paper is to find ways to treat the cancer tumors before they can reach an advanced stage devel-opmen.

**AMS Subject Classifications:** 92B05, 37C45.

## 1. Introduction

In this paper some deterministic and stochastic mathematical models are considered, which explain the interaction of the immune system and cancer.

Any treatment, which can improve the bodies own immune response is called the immunotherapy.

The treatment by cytotoxic chemotherapy helps to kill rapidly dividing cells, but it can harm normal tissues at the same time. The use of immunotherapy in conjuction with cytotoxic chemotherapy is known as bio chemotherapy. The bio chemotherapy treatment is obtained by using the immunological drug Interleukin-2. The Interleukin-2 at time t is denoted by I(t). The cells, which can not kill other cells are called naive T-cells. The number of naive t-cells at time t is denoted by *T_N_*(*t*). The naive t-cells will have the ability to kill other cells, if activated by antigen presenting cells. The most important type of antigen presenting cells are known as immature dendritic cells, immunogenic dendritic cells and to tolerogenic dendritic cells. All cells population are assumed to be antigen specific.

In section 2, we shall write the most recent mathematical models, which incorporate some different cells and the cytosine IL-2.

In section 3, we generalize the models in section 2 to stochastic dynamical systems.

## 2. deterministic models and cancer

There are various mathematical models of cancer and immune response [1–6]. As an example, let *φ*(*t*) be the population of the immunogenic dendritic cells at time t and *ψ*(*t*) be the Immature dendritic cells. It is assumed that the cells *φ*(*t*) are produced at a constant rate a by the Immature dendritic cells *ψ*(*t*). It also decays at a constant rate *ω*. This yields the following equation:

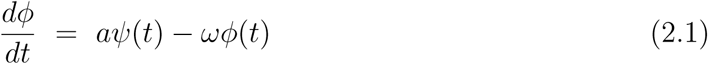

Let *M* (*t*) be the chemotherapy drug concentration in the blood stream at time t. The functions *M* (*t*) and *I*(*t*) decay exponentially with respect to *t*;

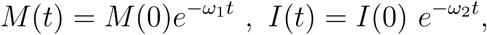

where *ω*_1_ and *ω*_2_ are positive constants.

The tumor cell population *T* (*t*) grows logistically in the absence of immune response, [7,8].

The function *T* (*t*) satisfies the equation

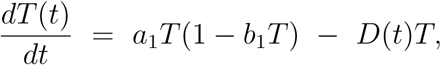

where

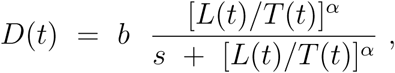

and *L*(*t*) is the number of cells, which combat a specific invader at time t.

Let *η*(*t*) be the number of tolerogenic dendritic cells. following [9,10,11], the function *η* has linear natural death term and satisfy the equation

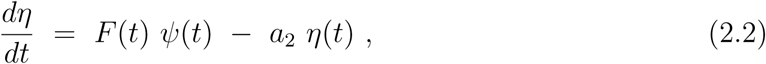

where F is given by

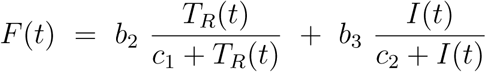

The function *T_R_* is the number of the so-called regularity T-cells at time t. The relation between *T_R_*(*t*) and *T_N_*(*t*) is given by:

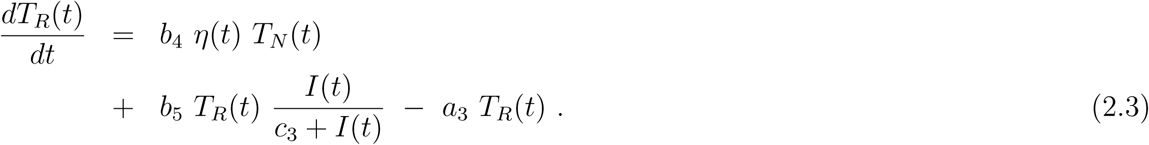

The function *T_N_*(*t*) satisfy the following equation;

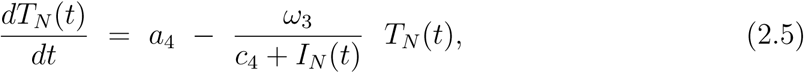

All the parameters *a, b*, *a*_1_, … *a*_4_*, b*_1_*, …, b*_5_*, c*_1_*, …, c*_4_ are nonnegative constants.

## 3. Stochastic models

It is preferred to find ways to treat tumors before they can reach an advanced stage of development.

Let us study the immunogenic dendritic cells *φ*(*t*).

To be more realistic, we consider the effect of the concentration of the chemotherapy M(t), and assume that *{φ*(*t*)*}* is a stochastic process satisfying a stochastic differential equation

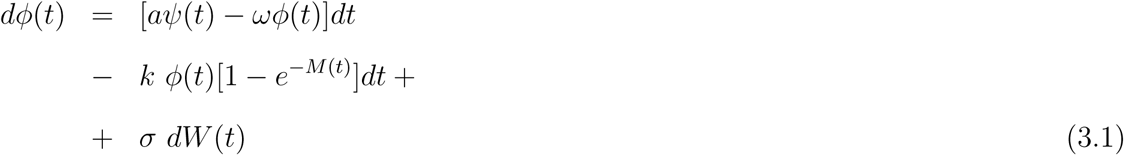

where *σ* is the volatility of the process {*φ*(*t*)} and W(t) is a Wiener process with zero mean and unit variance.

The function *ψ*(*t*) satisfies the equation

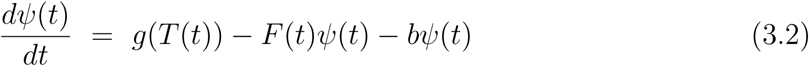

where *g* increases monotonically with respect to T(t), [8,9]. equation (3.1) can be written in the form

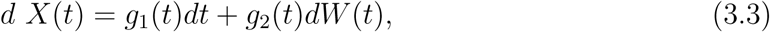

where

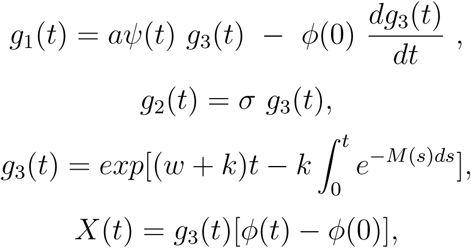

a, w, k and *α* are positive constants.

It is supposed that *φ*(0) is deterministic. It is clear that

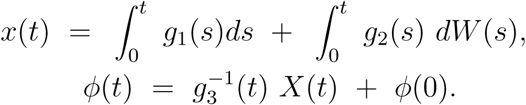

According to Feynman-Kac formula [9], we get

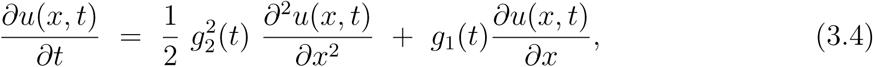

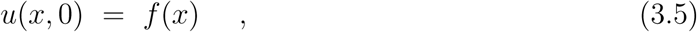

where

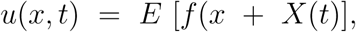

(E(Y) is the expectation of Y).

If f is continuous and bounded on (*−∞, ∞*), then the solution of (3.4), (3.5) is given by;

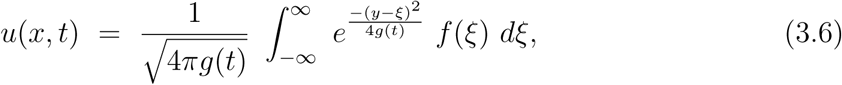

where

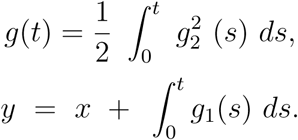

using (3.6), one gets that the characteristic function of the stochastic process *{X*(*t*)*}* is given by

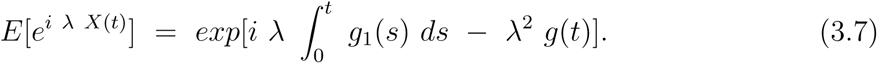

From (3.3), we get

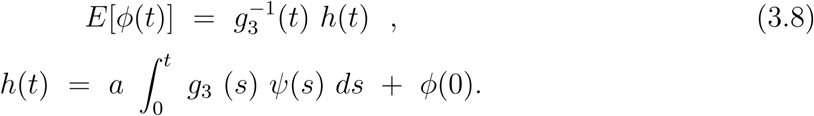

From (3.7), one gets

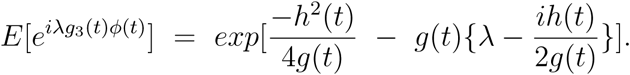

Using the last formula, we find that all the moments of the stochastic process {*φ*(*t*)} are given by:

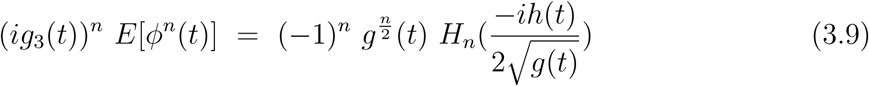

where *H_n_*(*x*) is the Hermite polynomial.

The variance of *φ* is given by 
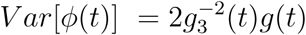
.

Our stochastic model,[10–16] should be viewed as an attempt to understand the growth dynamics of some dendritic cells and the effect of the chemotherapy.

In immunotherapy without IL-2 delivery, tumor-specific lymphocytes are reintroduced to the body after being inoculated with high concentrations of the cytokine IL-2. We add such treatment via immunotherapy to this model by shifting the initial lymphocyte population. (see [17–23]).

